# Silencing of ANKRD12 circRNA induces molecular and functional changes associated with invasive phenotypes

**DOI:** 10.1101/366245

**Authors:** Thasni Karedath, Ikhlak Ahmed, Wafa Al Ameri, Fatima M. Al-Dasim, Simeon S. Andrews, Samson Samuel, Iman K. Al-Azwani, Yasmin Ali Mohamoud, Arash Rafii, Joel A. Malek

**Author notes:** These authors contributed equally to this work (Thasni Karedath, Ikhlak Ahmed). Correspondence to: Joel A. Malek.

## Abstract

Circular RNAs (circRNA) that form through non-canonical backsplicing events of pre-mRNA transcripts are evolutionarily conserved and abundantly expressed across species. However, the functional relevance of circRNAs remains a topic of debate. In this study, we identified and characterized a circular RNA derived from Exon 2 and Exon 8 of the ANKRD12 gene, termed here as circANKRD12. We show that this circRNA is abundantly expressed in breast and ovarian cancers. The circANKRD12 is RNase R resistant and predominantly localized in the cytoplasm in contrast to its source gene mRNA. We confirmed the expression of this circRNA across a variety of cancer cell lines and provide evidence for its functional relevance through downstream regulation of several tumor invasion genes. We show that silencing of circANKRD12 induces a phenotypic change by significantly regulating cell cycle, increasing invasion and migration, altering the metabolism in cancer cells. These results reveal the functional significance of circANKRD12 and provide evidence of a regulatory role for this circRNA in cancer progression.

## 1. INTRODUCTION

Exonic Circular RNAs (circRNAs) were recently discovered as a new class of RNAs, functionally still largely uncharacterized. They possess distinct properties compared to linear RNAs and arise from direct backsplicing events that covalently link the 3’ end of an exon with the 5’ end of either the same exon or any other further upstream exon [1].

circRNAs were initially considered as molecular artifacts of aberrant RNA splicing [2]. This hypothesis was challenged by the observation that circRNAs are detected in various cell types in an evolutionarily conserved manner [3]. The copy number of circRNAs can be up to 10 times greater than that of associated linear RNAs, suggesting that these circRNAs may possess biological functions [3,4]. Studies have shown that some circRNAs harbor multiple binding sites for microRNAs, thereby "sponging” microRNAs and serving as competitive inhibitors for microRNA functions [5]. Some circRNAs have been shown to interact with RNA binding proteins to form RNA protein complexes thereby regulating canonical linear splicing of the gene [6]. These findings suggest that circRNAs hold dynamic and distinct roles in gene regulation.

Emerging evidence suggests that regulation of circRNAs is closely associated with different diseases, particularly cancer with aberrant expression pattern [7–10]. Thus, circRNAs represent a new class of diagnostic biomarkers with potential therapeutic significance [11,12]. The longer half-lives compared to their linear counterparts makes circRNAs long-acting regulators of cellular behavior and robust biomarkers [13,14]. A growing body of evidence has implicated functional involvement of circRNAs in regulating cancer progression and proliferation [8,15,16].

In the present study, we investigated the functional role of a high abundance circRNA from Ankyrin Repeat Domain 12 (ANKRD12) gene in cancer progression. ANKRD12 is a paralog of ANKRD11, a putative tumor suppressor gene with multiple functions including as a p53 co-activator [17,18]. Low ANKRD12 level is an independent prognostic predictor of colorectal carcinoma patients [19]. Circular isoforms of ANKRD12 have been identified in cancer cells and patient samples in many recent studies including ours [20,21]. In this study, we validated one of the most predominant circular isoform of ANKRD12 gene that includes the backsplice junction of exons 8 and 2 (circANKRD12). We report that circANKRD12 regulates the invasion, migration proliferation and cellular bioenergetics of cancer cells by modulating cell signaling, metabolic and cell cycle regulation pathways in cancer. We confirmed the expression of this circRNA across a variety of cancer cell lines and provide evidence for its functional relevance through downstream regulation of several tumor invasion genes.

## 2. MATERIALS AND METHODS

### 2.1. Cell Lines and treatment

Ovarian cancer cell lines PA1, SKOV3, CAOV3, OVCAR3, breast cancer cell lines MDA-MB-231, MCF7 and T47D and breast normal cell line MCF-10A, Lung cancer cell lines NCI-H226 and Lung Normal Fibroblast cell line LL24 (all from American type Cell Collection, Manassas, VA), APOCC (ovarian primary cell line derived from ascites fluid) (pers. communication Dr. Arash Tabrizi), A2780, A2780 CIS (Sigma, St. Louis, MO) were used for the current study. Cells were cultured in DMEM (Life technologies, NY, USA) supplemented with 10% fetal bovine serum (Life technologies, USA).

### 2.2. RNA preparation and qRT-PCR

The nuclear and cytoplasmic RNA were extracted using The SurePrep™ Nuclear or Cytoplasmic RNA Purification Kit (Fisher Grand Island, NY, USA). Total RNA from whole-cell lysates by using RNAesay mini kit (Qiagen Valencia, CA USA). For RNAse R treatment, 2 μg of total RNA was incubated 60 min at 40 °C with or without 3 U μg^-1^ of RNAse R (Epicentre Technologies, Madison, WI), and the resulting RNA was subsequently purified using an RNeasy MinElute cleaning Kit (Qiagen, Valencia, CA USA). cDNA synthesis was carried out using First strand synthesis kit (AMV) from Roche Biosciences or Biorad select cDNA synthesis kit using random primer for circRNA experiments. Fast Start Universal *SYBR Green* Master mix (Roche, Clovis, CA) was used to amplify the specific gene using cDNA primes obtained from Primer bank (http://pga.mgh.harvard.edu/primerbank/ (Supplementary Table 1). Each Real-Time assay was done in triplicate on Step One Plus Real time PCR machine (Life Technologies, CA, USA).

### 2.3. Transfection

siRNA transfection was carried out using custom designed siRNAs for both ANKRD12 circular and linear transcripts (Fig 2 and Supplementary Table 1). The SKOV3, MDA-MB-231, OVCAR3, NCI-H226 cells were grown in 6 well plates for transfection. The cells were transfected at 24h with 30pmol concentration of siRNA (VWR, Radnor, PA, USA) or scrambled control (Mission siRNA universal negative control, Sigma, St.Louis, USA) using Lipofectamine RNAi max (Invitrogen MA USA) according to manufacturer’s protocol. These experiments were conducted in three different biological triplicates for subsequent RNA-sequencing.

### 2.4. RNAseq Analysis

SKOV3, OVCAR3, NCI-H226, MDA-MB-231 cells transfected with cirANKRD12 or universal scrambled control were used for RNAseq analysis. RNA-seq library preparation and *In silico* detection of circRNA candidates from paired end RNA-seq data was conducted as described earlier [20]

### 2.5. Cell proliferation assay

Cells were cultured at a density of 5×10^3^ cells per well in flat-bottomed 96-well plates and supplemented with 10% DMEM with antimycotic antibiotic. Experiment was done at different time points (24 h and 48 h or based on cell doubling time). CellTiter 96^®^ Aq_ueous_ One Solution Reagent (Promega, Madison, WI) was added and experiment was conducted according to the manufacturer’s instructions.

CellTiter-Glo assay (Promega Madison USA): experiment was conducted according to the manufacturer’s instructions. Briefly, CellTiter-Glo reagent was added directly to the wells of 96 well plate and luminescence was measured on an Envision reader (PerkinElmer).

### 2.6. Scratch assay- cell migration assay

The scratch assay was conducted as described earlier [22]. Cells were plated into a 6-well plate with complete medium and grown to 80% confluence. Cells were transfected with respective siRNA in OPTIMEM medium, and the medium was replaced with serum free DMEM after transfection. After cells were grown to 100% confluence, a wound was created by scraping the confluent monolayer cells with a p200 pipette tip. Cells were then grown either in serum free medium or medium containing 3mM Thymidine. The distance between the two sides of cell-free area was photographed using 10x objective in an AXION Zeiss epiflurescence microscope. The distance is measured using Zeiss Zen software (Carl Zeiss Carpenteria, CA, USA,)

### 2.7. Trans-well migration and invasion assay

Cellular migration and invasion was determined using a Trans well Boyden chamber assay as described previously [23].

### 2.8. 3D organotypic spheroid Model experiments

3D anchorage independent spheroids were developed in SKOV3 cancer cell lines; initially cells were seeded on ultra-low attachment plate (Corning, NY, USA) for 3 days to facilitate spheroid formation. Transfection of the spheroids with circANKRD12 siRNA was conducted using the reverse transfection method.

### 2.9. Spheroid area measurement

Cells were seeded at a density of 300 000 cells/well into ultralow attachment plats. After 72h, cells were transfected either with scrambled siRNA or circANKRD12 siRNA. After 48h the diameters of at least 50 spheroids were measured and spheroid area was measured Zeiss Zen Software.

### 2.10. Cell Proliferation assay in 3D organotypic models

10000 cells were seeded on each well of ultra-low attachment 96 well plate with OPTIMEM medium. After three days, once spheroids were formed, transfection was conducted with circANKRD12 sirRNA or scrambled control. After 24h and 48h of transfection, MTS reagent was added to the medium. Measurements were according to manufacturer’s instructions.

### 2.11. Collagen Invasion assay in 3D organotypic models

3D organotypic models of either circANKRD12 transfected or scrambled control SKOV3 cells expression GFP were placed of the top of jellified collagen matrix (Rat tail collagen1, 1mg/ml). The invasiveness analyzed after 24h to 10 days.

### 2.12. Cell Cycle Analysis

Cell cycle analysis was done on cells fixed with FxCycle™ PI/RNase Staining Solution using BD LSRFortessa™ cell analyzer.

### 2.13. Western Blot Analysis

Cellular protein was extracted after 48h of transfection. The cells were lysed in 100ul of RIPA buffer with protease inhibitor cocktail. Then 40 micrograms of protein were resolved in SDS PAGE gel and transferred to a nitro cellulose membrane. The primary antibodies used were anti Cyclin D1, Anti Cylin B1, Anti CyclinD2, anti-Cyclin Phospho B1 and β-actin (cell Signaling, USA). The blots were visualized by ECL detection (Amersham, N J, United States,).

### 2.14. circANKRD12 and ANKRD12 siRNA longevity assay

siRNA longevity was calculated based on cell doubling time (10^th^ doubling time [20days]). Cell doubling time of SKOV3 cells was calculated based on the mathematical equation.

DoublingTime=duration*log (2) / log(FinalConcentration)-log(InitalConcentration). Cells were seeded on 60mm petri plates and transfection was done with siRNA of circANKRD12, or ANKRD12 mRNA. Cells were harvested on an interval of 48h (doubling Time). Experiment was extended till 10^th^ doubling time (20days). The silencing efficacy was measured by gene expression analysis (RT-PCR).

### 2.15. Assessment of mitochondrial function by seahorse extracellular flux analyzer

The mitochondrial oxygen consumption rate (OCR) in the MDA-MB-231 and SKOV3 cells was assessed by using a Seahorse Bioscience XFe96 analyzer (Massachusetts, USA). Seahorse Bioscience XF Cell Mito Stress Test assay kit was used for the study [22]. Experiments were conducted as per manufactures protocol. 10000 cells were seeded on assay plate and assay was completed in 48 hours after the transfection.

### 2.16. Statistical Analysis

Statistical analysis in all the experiments is based on at least three biological replicates and the error bars are drawn with standard error of means (SEM). The p value is calculated by using student’s T test.

## 3. RESULTS

### 3.1. Characterization of circANKRD12 in cancer cells

We characterized the circular RNA expression using RNA sequencing (RNA-seq) of total RNA in ovarian cancer cell lines (SKOV3 and CAOV3) undergoing Epithelial to Mesenchymal Transition. Using our circRNA detection pipeline [20], we identified a total of ~18,700 circRNAs in these cell lines. The circRNA from ANKRD12 gene having a backsplice junction between exons 8 and 2 (circANKRD12) was one of the highly abundant circular RNAs in both SKOV3 and CAOV3 cell lines. The circANKRD12 originates from 18p11.22 chromosome locus with the backsplice junction forming between exons located ~39.3kb apart (Fig 1a). By carefully designing two sets of divergent primer pairs in Exon2 and Exon8 and sanger sequencing of the amplified products, we were able to identify two circular isoforms of 286bp and 925bp lengths sharing the same backsplice junction (Supplementary File 2-S2:1-3), Supplementary Table 2). The first isoform of 286bp is comprised of only two exons (Exon2 and Exon8), while the 925bp isoform is comprised of 6 exons (Exons 2,3,5,6,7 and 8). All subsequent functional studies do not distinguish between these two circular isoforms as they targeted the junction sequence that is unique to circular forms vis-à-vis linear forms.

**Fig 1.**
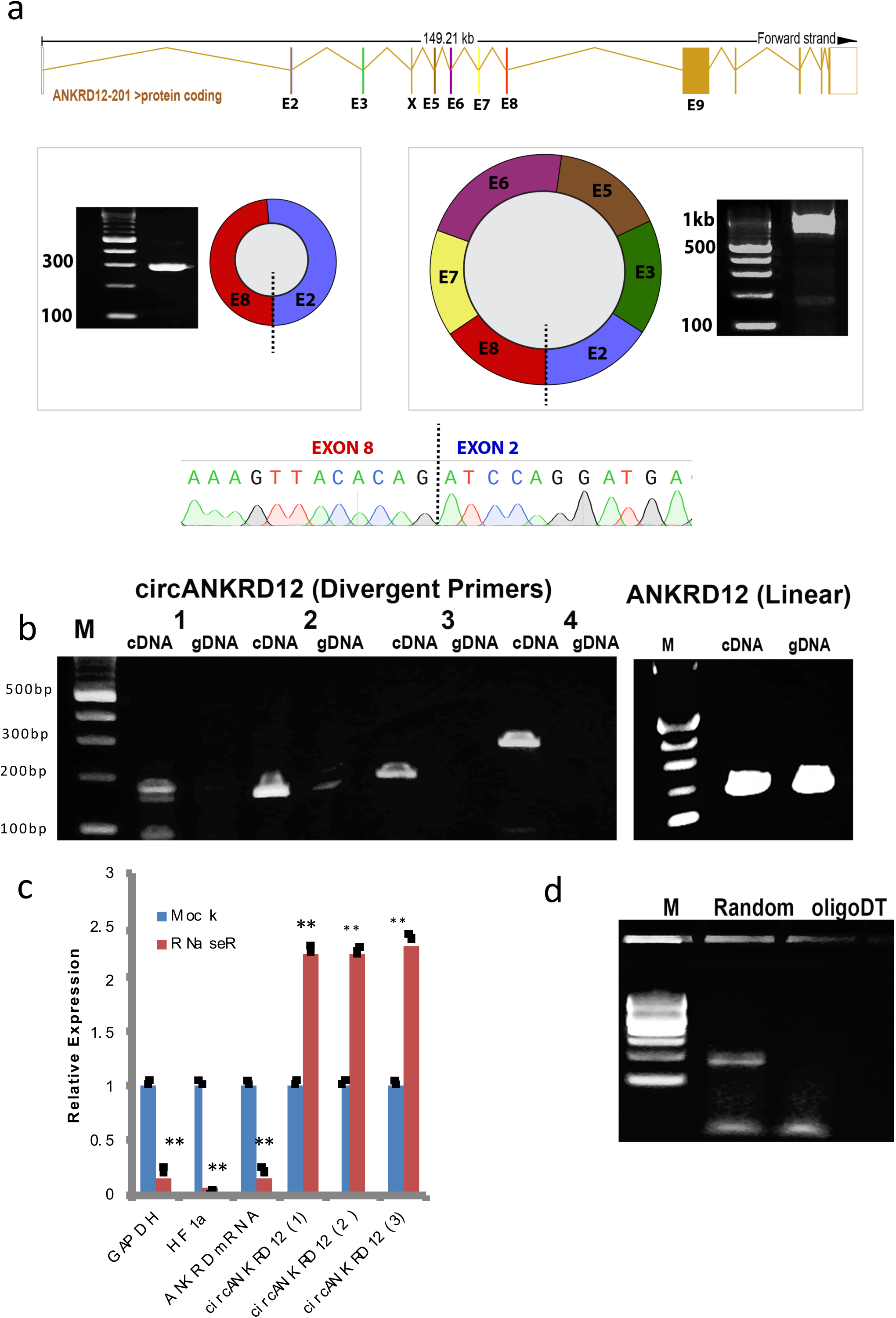
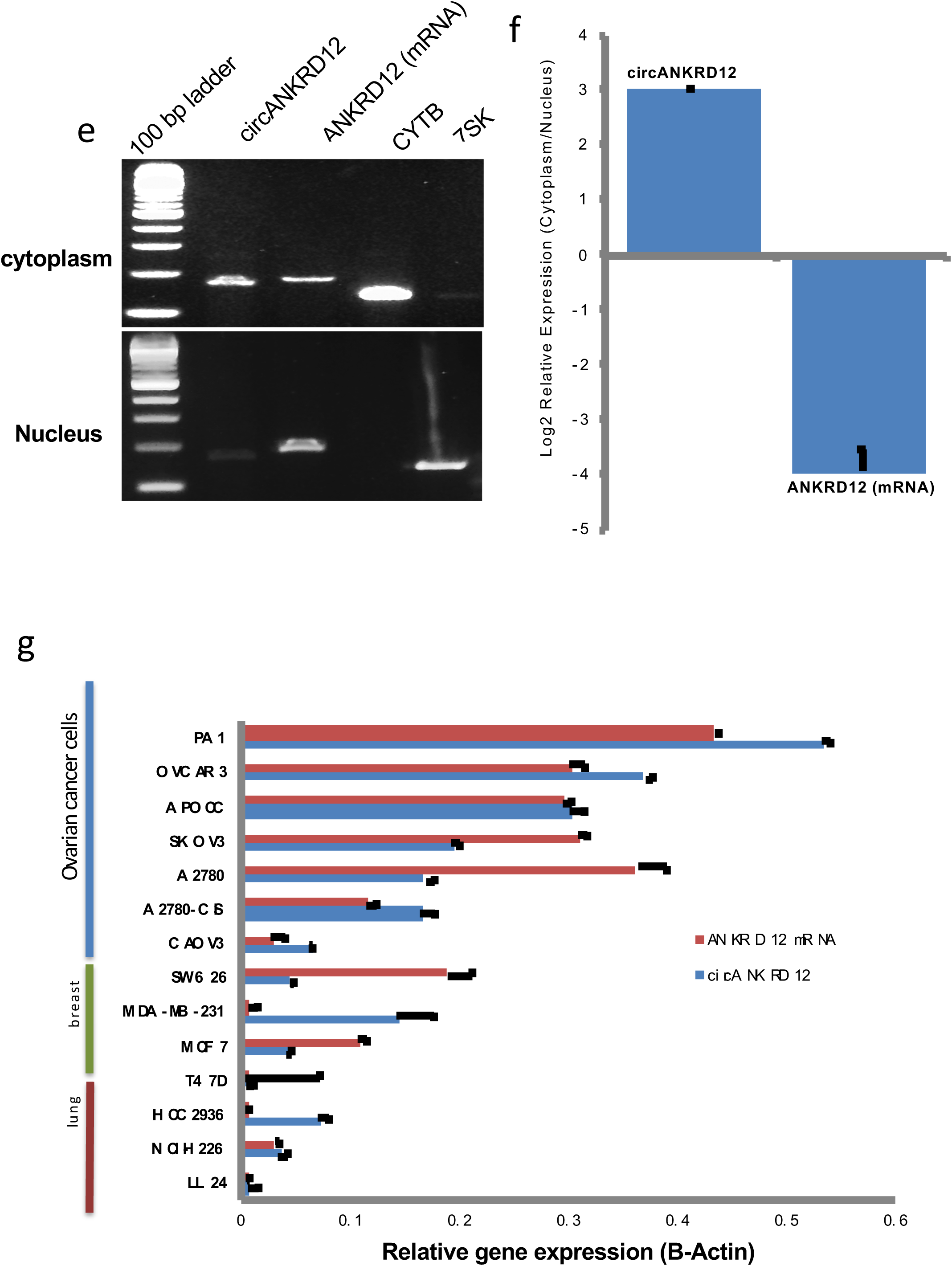
Characterization of circANKRD12 in human cancer cells. **(a)** Transcript structure of the ANKRD12 gene. Two predominant circular isoforms (286 bp and 925 bp) were detected sharing the same backsplice junction between Exon8 and Exon2 as represented by the dashed line. The backsplice junction was sequenced for validation. **(b)** Divergent primers designed to detect the circANKRD12 backsplice junction amplified cDNA but not genomic DNA (4 pairs of different primers were used to amplify the circular junction). Linear ANKRD12 amplicon was detected in both cDNA and gDNA. M represents 100 bp ladder. **(c)** qRT-PCR for relative abundance of circANKRD12 and linear ANKRD12, GAPDH and HIF1 alpha in SKOV3 cells treated with RNase R **(d)** Backsplice junction was detected in cDNA synthesized using random hexamer but not oligoDT primed RNA. **(e)** Abundance of circANKRD12 and ANKRD12 linear RNA. Semi qRT-PCR data indicating abundance of circANKRD12 and ANKRD12 mRNA in the cytoplasmic and nuclear fractions of SKOV3 cells, nuclear and cytoplasmic purity markers were also assessed using nuclear specific marker 7SK and cytoplasmic specific marker CYTB. The gel picture represents PCR amplified products of circular RNAs with three different sets of primers. **(f)** Real time data shows the abundance of circANKRD12 and ANKRD12 mRNA in cytoplasm and nucleus. (h) Abundance of circANKRD12 and ANKRD12 mRNA in a panel of cancer cells. The data in **c,g and h** are means with error bars representing standard error of mean (SEM) from three experiments; \*\**P*<0.01 (Student’s t-test).

Multiple validation experiments were used to confirm circANKRD12 expression (Fig 1 b-d). Divergent primers were designed to amplify the backsplice exon junction. As expected, each primer pair produced a single distinct band of expected product size in RT-PCR assay indicating the presence of the circular junction. The backsplice junctional sequence was confirmed by Sanger sequencing. The divergent primers, with respect to genomic sequence, only amplified when cDNA was used as a template. The same primers did not produce a product from genomic DNA (gDNA). Conversely, PCR using convergent primers with respect to genomic sequence could amplify both cDNA and gDNA templates from ANKRD12 gene (Fig 1b). This strongly indicates that a head-to-tail backsplicing junction in circANKRD12 only exists in the RNA form.

To examine whether circANKRD12 is resistant to exonucleases, we treated the total RNA extracted from the SKOV3 cell line with RNase R. This exoribonuclease enzyme digests all linear RNA forms with a 3’ single stranded region of greater than 7 nucleotides [24]. As circRNAs are devoid of any 3’ single strand overhangs, they are expected to show resistance to digestion by RNase R. Indeed, circANKRD12 was resistant to RNase R digestion compared to linear forms of ANKRD12, HIF1 alpha and GAPDH. Resistance to digestion with RNase R exonuclease confirmed that the circANKRD12 is a stable circularized transcript (Fig 1c). Further, cDNA created by priming with oligo (dT) primers failed to produce any PCR amplification products for circANKRD12. On the other hand, priming with random hexamers, resulted in distinct PCR bands for the backsplice junction shows circANKRD12 is not poly A tailed (Fig 1d). The RT-PCR analysis of nuclear and cytoplasmic fractions of RNA demonstrated that ANKRD12 circRNA is predominantly localized in the cytoplasm (Fig 1e,f). The purity of cytoplasmic and nuclear fractions was confirmed by amplifying the fractions using nuclear and cytoplasmic specific markers supplementary File 2 (S2:8)

We performed Real-time PCR analysis on 12 different cell lines from breast, ovarian and lung cancer and normal breast and lung to assess the cell-type specific expression of circANKRD12 (Fig 1g). Most of these cell lines show a high abundance of both ANKRD12 circRNA and mRNA. Ovarian cancer cell lines show a higher abundance of ANKRD12 circRNA compared to breast and lung cancer cell lines.

### 3.2. siRNA-mediated knockdown of circANKRD12 is highly specific

To investigate the role of circANKRD12, we designed siRNAs to target the backsplice junction (Fig 2a) and transfected multiple cancer cell lines to induce siRNA-mediated knockdown of the circle while leaving the linear RNA unaffected. The circRNA specific siRNA was designed against the backsplice junction spanning exons 2 and 8 of the gene. We observed high knockdown efficiency of greater than 90% of the circular junction when using the siRNA versus the control (scrambled siRNA) in the four cell-lines (Fig 2b). Using two siRNA constructs against the circANKRD12, we confirmed that knockdown of the circular RNA is specific and has no significant effect on the linear mRNA expression (Fig 2c). We also designed siRNAs targeting exon9 and exon7 of ANKRD12 gene to knockdown the linear mRNA. Indeed, the siRNA designed against exon 9, which lies outside the circANKRD12 locus was successful at knocking down the mRNA and exhibits no effect on circANKRD12 levels (Fig 2d). We investigated the specificity of siRNA constructs designed against circANKRD12, Exon7 and Exon9 through a series of knockdown experiments explained in supplementary File 2 (S2:4-7).

**Fig 2.**
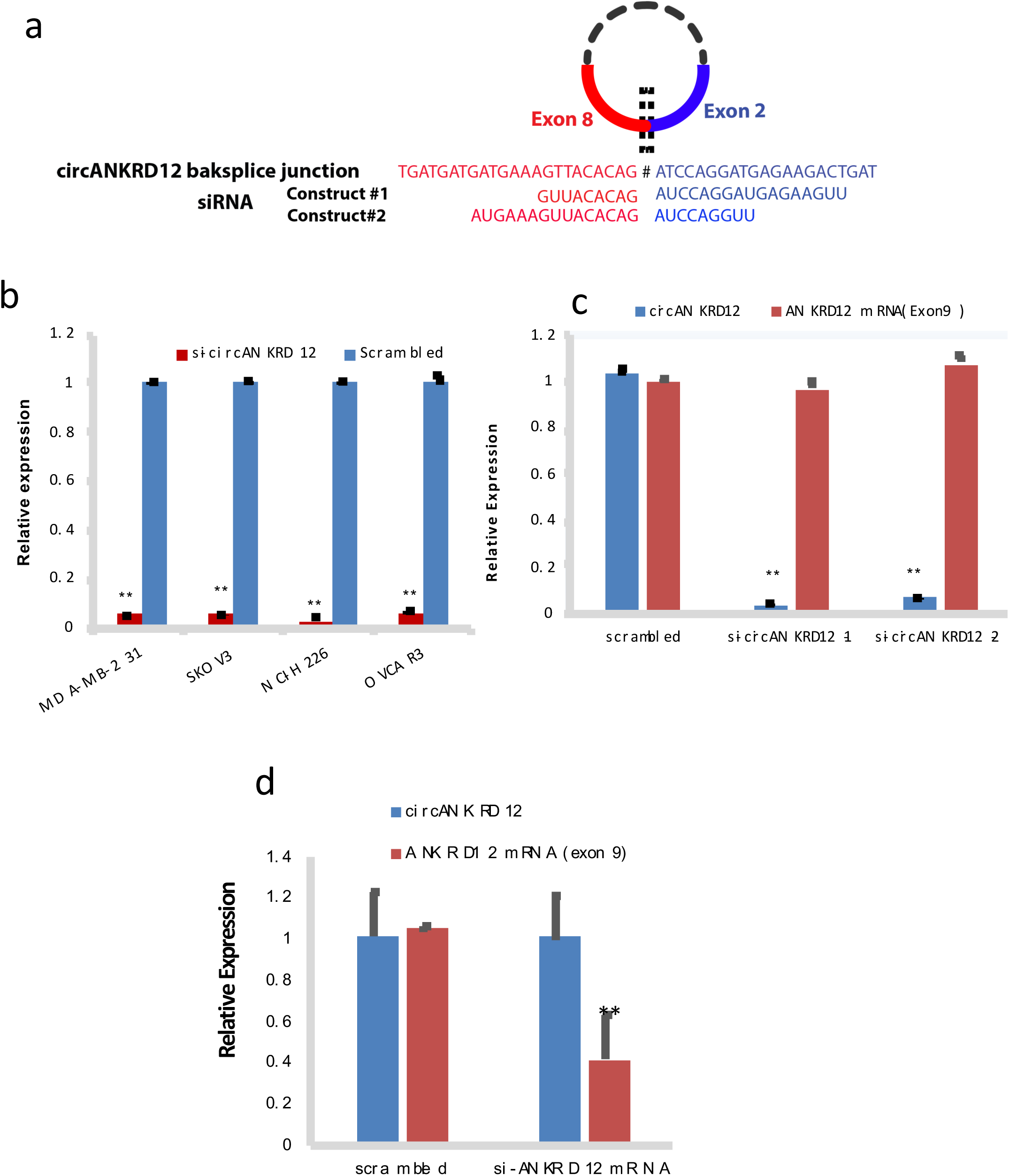
siRNA mediated silencing of circANKRD12 in cancer cells. (**a**) Two circANKRD12 siRNAs spanning the back-splice junction (**b**) qRT-PCR analysis for knockdown efficiency of circANKRD12 siRNA in 4 different cell lines. (**c**) qRT-PCR analysis for knockdown efficiency of two different circANKRD12 siRNA constructs in SKOV3 cells. (**d**) qRT-PCR analysis for silencing efficiency of ANKRD12 linear siRNA (exon9) in SKOV3 cells. (Data in **b**-**d** are the means with error bars indicating standard error of mean (SEM) of three experiments. \*\**P* < 0.01 (Student’s *t*-test)

### 3.3. Silencing of circANKRD12 changes molecular phenotypes of ovarian, breast and lung cancer cells

RNA-sequencing was performed in triplicate for two ovarian cancer cell lines (SKOV3, OVCAR3), breast (MDA-MB-231) and a lung cancer cell line (NCI-H226) for both scrambled control and siRNA targeting circANKRD12. **Table 1** gives the number of differentially expressed genes in circANKRD12 knockdown samples (Supplementary File S1) at a false discovery rate of less than 5% with at least 1.5-fold changes in expression.

**Table 1:** Number differentially up and downregulated genes for four cancer cell lines in siRNA-mediated knockdown of circANKRD12

|  | **Si-circANKRD12 vs. Scrambled** |  |  |  |
| --- | --- | --- | --- | --- |
|  | SKOV3 | OVCAR3 | MDA-MB-231 | NCI-H226 |
| Upregulated genes | 25 | 104 | 237 | 168 |
| Downregulated genes | 9 | 71 | 179 | 3 |

The canonical pathways analysis by Ingenuity’s IPA toolkit (IPA^®^, QIAGEN Redwood City, www.qiagen.com/ingenuity) revealed an overwhelming enrichment of differentially regulated genes for cell cycle regulation, tumor cell invasion, migration interferon signaling pathways and immune response of cells (Fig 4, supplementary Table 3). We observed an upregulation of inflammatory pathways such as tumor necrosis pathway, NFkB pathway, interferon signaling and down regulation of the cell cycle pathway (Supplementary File 2-S2:15). Silencing of circANKRD12 affects cellular bioenergetics by downregulation of key genes involved in oxidative phosphorylation, AMPK pathway and cell metabolism (Supplementary Table 4). IPA predicts a significant activation of interferon signaling pathway through upregulation of STAT1, STAT2, IFT1, MX1genes, and downregulation of estrogen mediated S phase entry through downregulating cyclin D1 and cyclin A. Activation of IL8 signaling and inflamasome pathways, closely associated with immune modulation is also predicted. IL8 signaling pathway upregulates IL8, IRAK, ICAM-1, COX-2 and VEGF genes there by upregulating angiogenesis, inflammation, and inhibits cell proliferation by downregulating cyclin D1. The differentially regulated genes cyclin D1 (CCND1), CBX5, STAU1 and AK4 were found to be consistently downregulated in the circANKRD12 knockdown across multiple cell lines (Fig 3a). Using qRT-PCR, we validated the differential expression of few selected genes in SKOV3 cells (Fig 3c). The gene cyclin D1 (CCND1) was among the genes that showed consistent downregulation in the circANKRD12 knockdown across multiple cell lines (Fig 3a). To ensure that the effect on CCND1 is related to circANKRD12 knockdown rather than off-target effects we compared the knockdown effects in SKOV3 (High in circANKRD12 and high in CCDN1) and LL24 cells (low in circANKRD12 and high in CCDN1) (Fig 3b). Unlike SKOV3 cells, the circANKRD12 level is low in LL24 cell line, attempts at knocking down circANKRD12 should not affect the cyclin D1 level in the cell line. Indeed, transfection of the siRNA against circANKRD12 in LL24 showed reduction of the already low levels of circANKRD12 but no effect on cyclin D1 expression, suggesting the siRNA effect on CCND1 is likely mediated through circANKRD12. Because cell cycle regulation seems to be one of the most deregulated pathways in circANKRD12 silenced cells, we decided to follow it up with further functional screening using cell based phenotypic assays.

**Fig 3.**
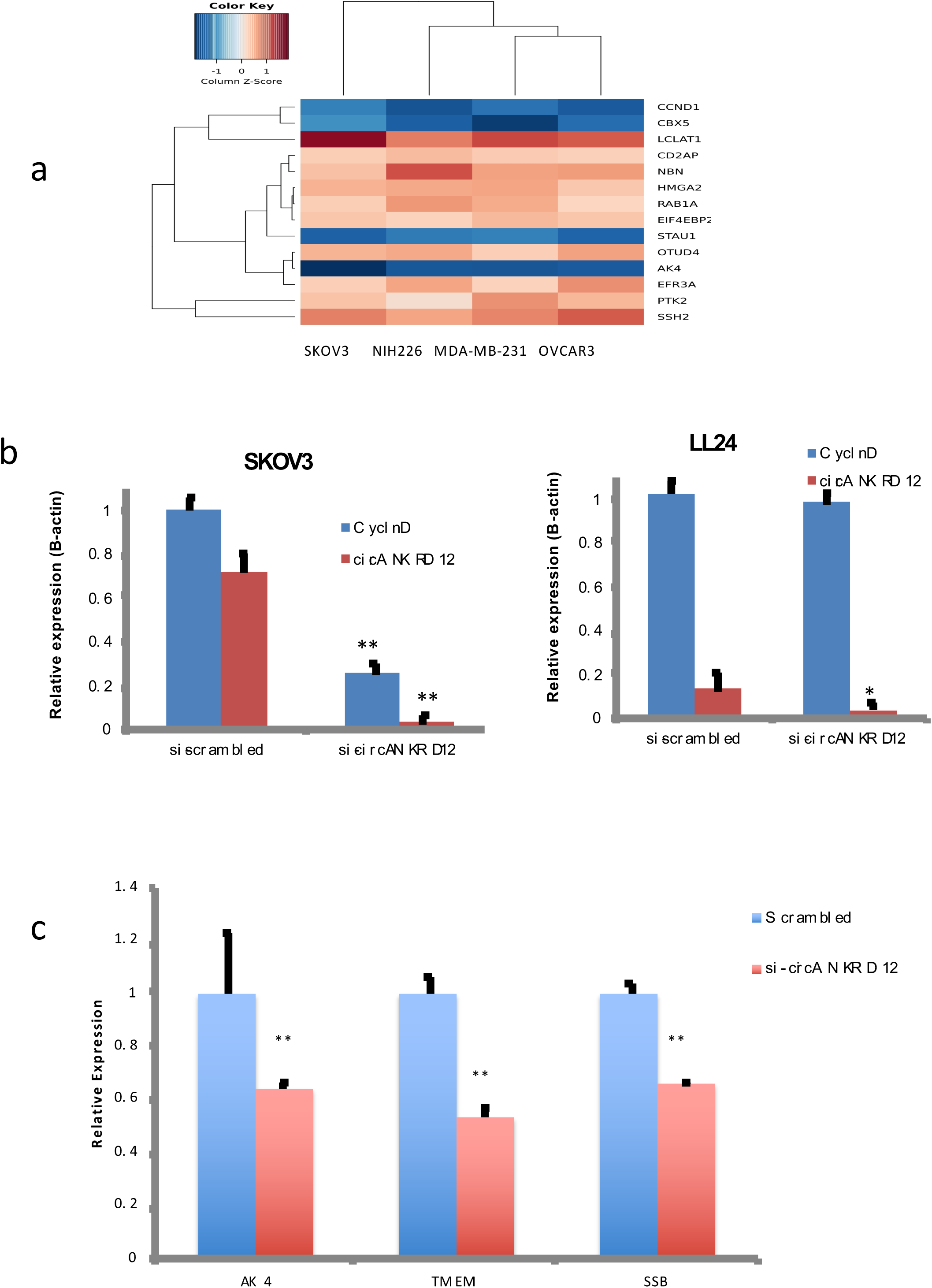
Silencing of circANKRD12 affects gene expression changes in cancer cells. (**a**) Heat map of differentially expressed genes from RNAseq analysis of SKOV3, NCI-H226, MDAMB 231 and OVCAR3 cells silenced with si-circANKRD12 compared to scrambled control. (**b**) qRT-PCR validation of expression level of cyclin D1 in circANKRD12 silenced SKOV3 and LL24 cell lines. In SKOV3 cells, circANKRD12 knockdown leads to a significant downregulation of Cyclin D1 (P=7.8E-04), while as in LL24 Cyclin D1 shows a minimal non-significant change in expression (P=0.18). (**c)** qRT-PCR validation of some selected genes differentially expressed in SKOV3 cells silenced with circANKRD12. Data in **b-c** are the means with error bars indicating standard error of mean of three experiments. \*\**P* < 0.01 (Student’s *t*-test).

**Fig 4.**
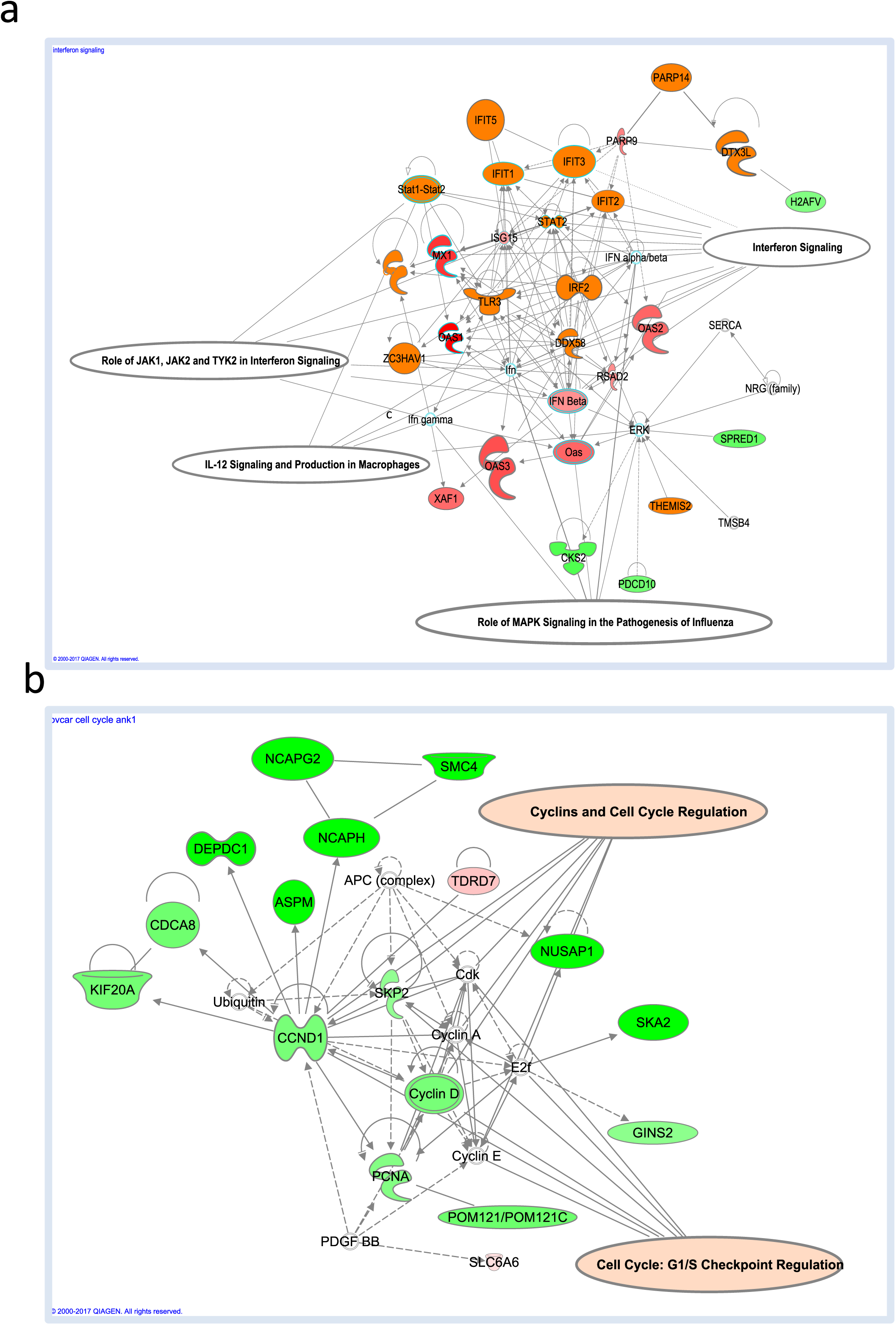
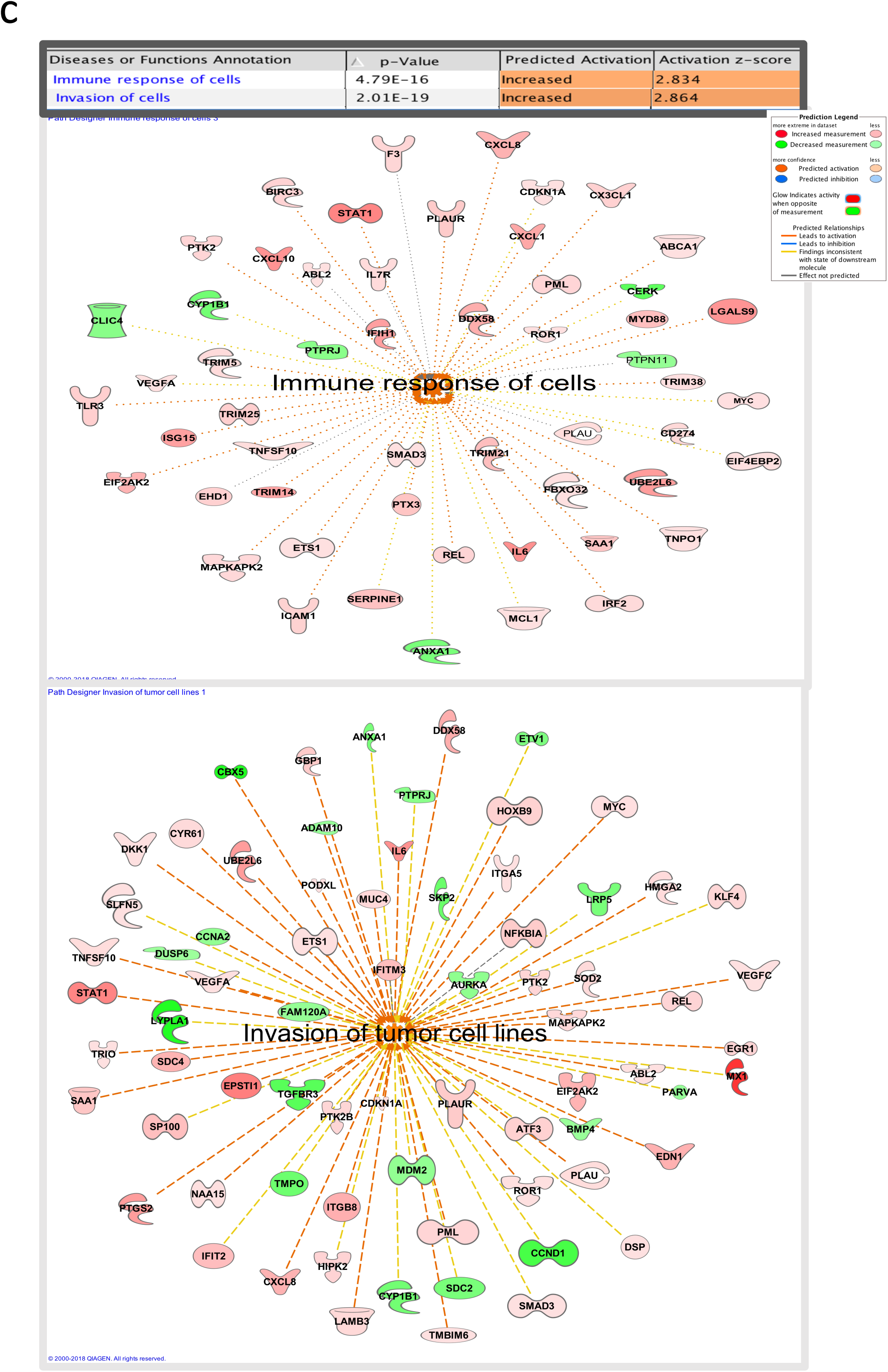
Gene network analysis of RNA-Seq data from MDA-MB-231 and OVCAR3 cells. (a) Gene network analysis using ingenuity pathway analysis in MDA-MB-231 cells silenced for circANKRD12 shows upregulation of Interferon, JAK/STAT, IL-12 signaling pathways (b) Gene network analysis using ingenuity pathway analysis in OVCAR3 cells silenced with circANKRD12 shows downregulation of Cyclins and Cell Cycle regulation pathways. (C) Shows functional predicted networks from RNAseq data of circANKRD12 silenced MDA-MB-231 cells induces inflammatory immune responses and cancer cell invasion with activation Z score and p values.

### 3.4. Abrogation of circANKRD12 by siRNA mediated silencing increases migration and invasion in ovarian cancer cells

Wound healing assays show that silencing of cirANKRD12 increased cell migration (Fig 5a b, Supplementary File2- S2:10), in SKOV3 cells after 24h. compared to scrambled control. This is confirmed again by cell migration assays in synchronized (Thymidine incorporation) cells, which show an increased migration rate for circANKRD12 silenced SKOV3 cells (Fig 5c). Matrigel invasion (inserts coated with matrigel) and migration analysis using Boyden chamber shows circANKRD12 silenced cells undergo significant increase in the invasion and migration compared to scrambled control (Supplementary File 2- S2:10).

**Fig 5.**
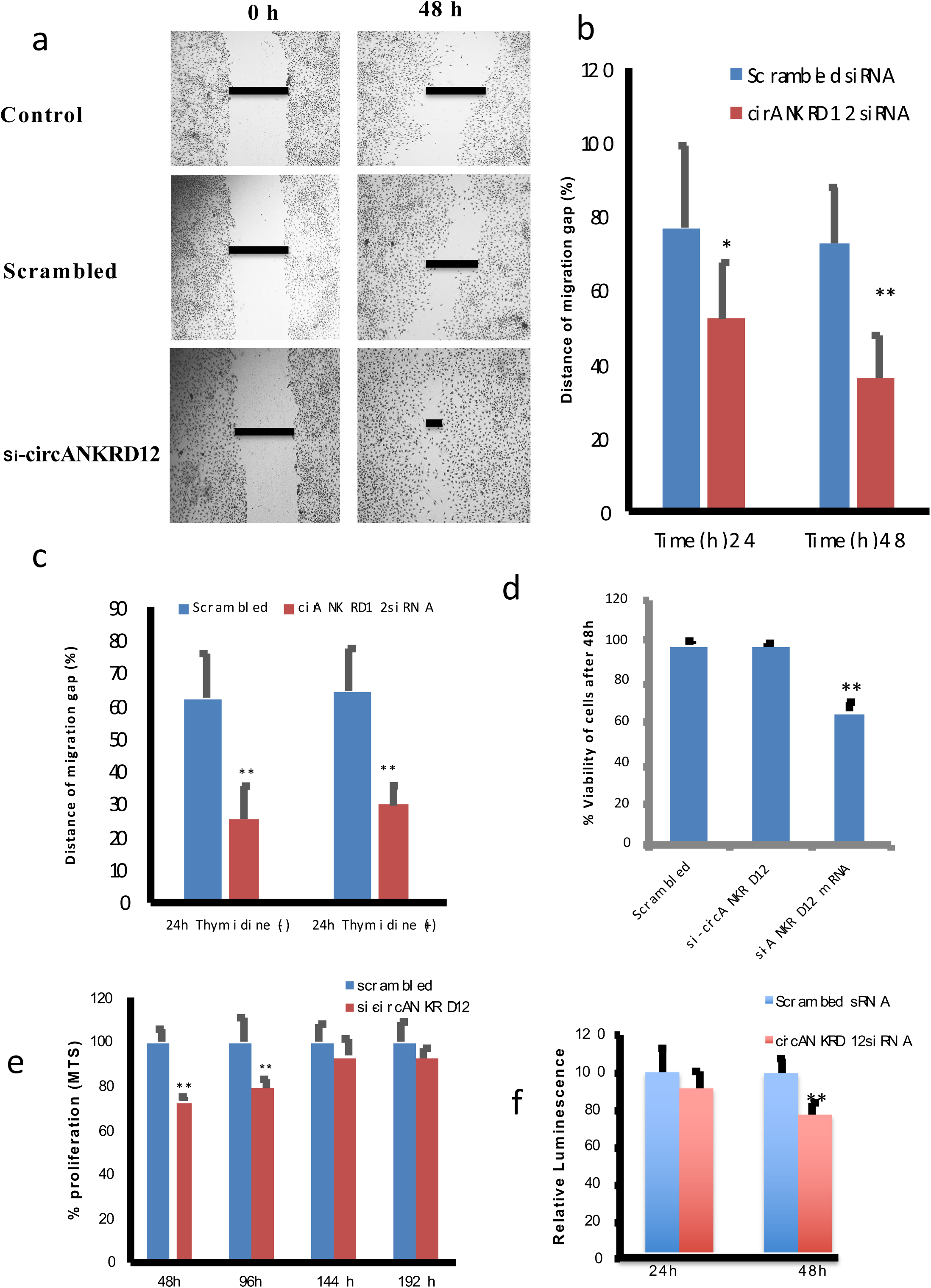
Silencing of circANKRD12 results in an invasive phenotype with low proliferation rate. (**a**) Representative Wound healing migration assay image of circANKRD12 silenced SKOV3 cells and its scrambled control at 0h and 24h (**b**) Bar diagram depicting the reduction in wound width after migration in scrambled and circANKRD12 silenced cells (average of 15 wounds from three different biological replicates). (**c**) Migration of circANKRD12 silenced SKOV3 cells after cell cycle synchronization with thymidine for 24h. (**d**) Cell viability is measured by using trypan blue exclusion assay. In circANKRD12 knockdown, the viability is reduced by roughly 2% while it is reduced by ~34% in ANKRD12-mRNA knockdown (**e**) Proliferation of SKOV3 cells transfected with siRNA against circANKRD12 and assessed using MTS cell proliferation assay kit at the indicated hours and data shown as % proliferation compared to control. (**f**) Cell titer glow assay shows relative luminescence rate of ATP production in circANKRD12 silenced SKOV3 cells compared to scrambled control. Data in **b**-**f** are the means with error bars indicating standard error of mean of three experiments. \*\**P* < 0.01 (Student’s *t*-test).

### 3.5. Silencing of circANKRD12 decreases cell proliferation

Knockdown of circANKRD12 and ANKRD12 mRNA significantly reduces cell proliferation (Fig 5, Supplementary File 2- S2:9). The results of MTS and ATP assays show a significant reduction in cell proliferation in circANKRD12 silenced cells compared to scrambled control (Fig 5e, f). The Trypan blue exclusion assays confirm the silencing of circANKRD12 is not affecting the cell viability significantly. However, silencing of ANKRD12 mRNA significantly reduces both proliferation and cell viability in SKOV3 cells (Fig 5d). These results indicate that knockdowns of both circular and linear RNA forms of ANKRD12 gene are capable of inducing strong phenotypic changes and modulate the growth or survival of cancer cells.

### 3.6. Silencing of cirANKRD12 in 3D tumor models induces a phenotypic switch from highly proliferative to invasive phenotype

Since 3D culture is known to have better cell-to-cell interactions and more closely mimics tumors *in vivo* [25], we investigated the effects of circANKRD12 knockdown in 3D culture experiments. The 3D anchorage independent growth of siRNA transfected SKOV3 cells showed smaller, dense aggregates associated with an invasive phenotype (Supplementary File 2-S2:12). This reduction in spheroid size in circANKRD12-silenced SKOV3 organotypic models results in an alteration of phenotype from less invasive to a more invasive one (Fig 6b). Cell proliferation MTS assay reveals a significant reduction in cell growth within a time period of 24 hours and 48 hours after knockdown of ANKRD12 circRNA (Fig 6c). Collagen invasion of the circANKRD12 silenced spheroids results in an increased invasion through the collagen gel with highly motile cells after 10 days of transfection (Fig 6d). These results indicate the phenotypic switching from a highly proliferative phenotype to a less proliferative phenotype with high invasion potential.

**Fig 6.**
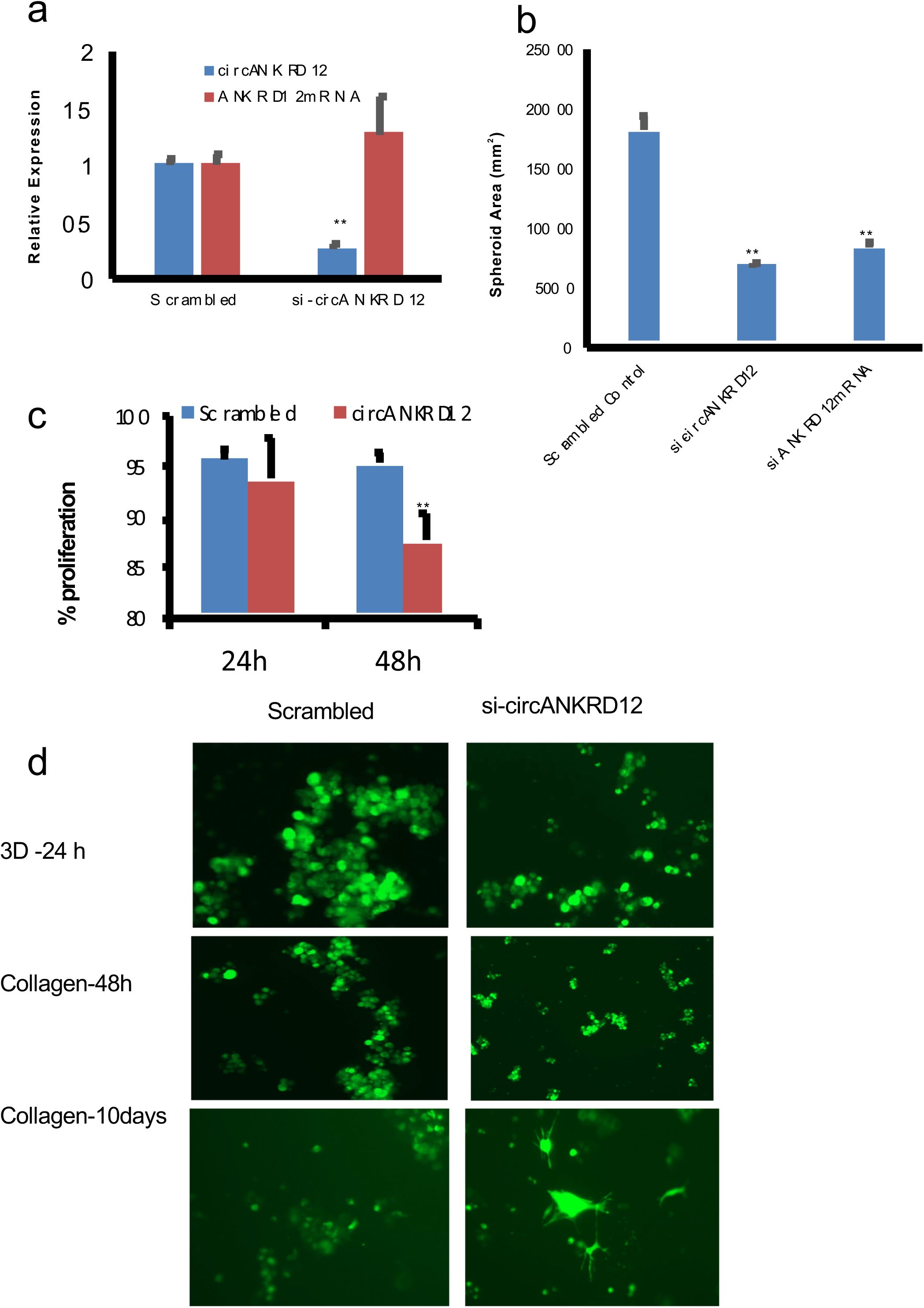
Silencing of circANKRD12 in 3D spheroid models of SKOV3 cells shows an invasive phenotype. (**a**) Knockdown efficacy of circANKRD12 in 3D cultures. siRNA directed against circANKRD12 downregulates circular RNA but shows no downregulation on mRNA of ANKRD12 gene in SKOV3 organotypic models (**b**) spheroid area is measured in anchorage independent 3D spheroids of SKOV3 cell lines, average of 100 spheroids in 3 independent experiments were measured. (**c**) MTS assay shows cell viability in 3D organotypic models is reduced at 24 and 48 h of transfection. (**d**) Collagen invasion assay of spheroids 24,48h and 10days. circANKRD12 silenced cells were able to invade through the thick collagen matrix and circANKRD12 silenced cells shows an invasive phenotype. Data in **a**-**c** are the means with error bars representing standard error of mean of three experiments. \*\**P*<0.01 (Student’s *t*-test)

### 3.7. Cyclin D1 is regulated in circANKRD12 silenced cells and involved in phenotypic switching by facilitating G1 arrest

circANKRD12 silencing results in cyclin D1 regulation and subsequent invasion and reduction in proliferation in ovarian cancer SKOV3 cells. Both real-time PCR and RNA-sequencing exhibited down regulation of cyclin D1 in circANKRD12 silenced cells. qRT-PCR analysis shows cyclin D1 expression is down regulated at least until 48 hours after transfection with the siRNA (Fig 7a). This is also supported by the western blot analysis, which shows a reduction in cyclin D1 protein level (Fig 7b). Only cyclin D1 shows differential regulation at the protein level in circANKRD12 silenced cells compared to other cyclin members including cyclin E1, pB1 and D2 (supplementary File 2-S2:13). To estimate the duration of the RNAi effect initiated by transfecting siRNA, we performed a longevity assay of circANKRD12 siRNA (si-circANKRD12). The longevity, initially stable for 6 days (supplementary File 2- S2:14) was checked on a cell doubling time basis estimated to be 48 hours (see Methods). The si-circANKRD12 exhibits an extended longevity period in SKOV3 cells, approaching 9 doublings (18 days). The reduction in the level of cyclin D1 is on par with the observed silencing efficiency and longevity of siRNA (Fig 7c). After the 10^th^ doubling, the effect on cell proliferation induced by silencing of circANKRD12 becomes insignificant (Fig 7d), suggesting that the knockdown efficiency is lost due to multiple passages of cells. As cyclin D1 is involved in cell cycle progression [26], cell cycle analysis of circANKRD12 knockdowns was conducted. Cell cycle analysis using FACS indicate there is a significant G0/G1 cell cycle arrest in SKOV3 cells silenced with circANKRD12 compared to scrambled control (Fig 8a, b).

**Fig 7.**
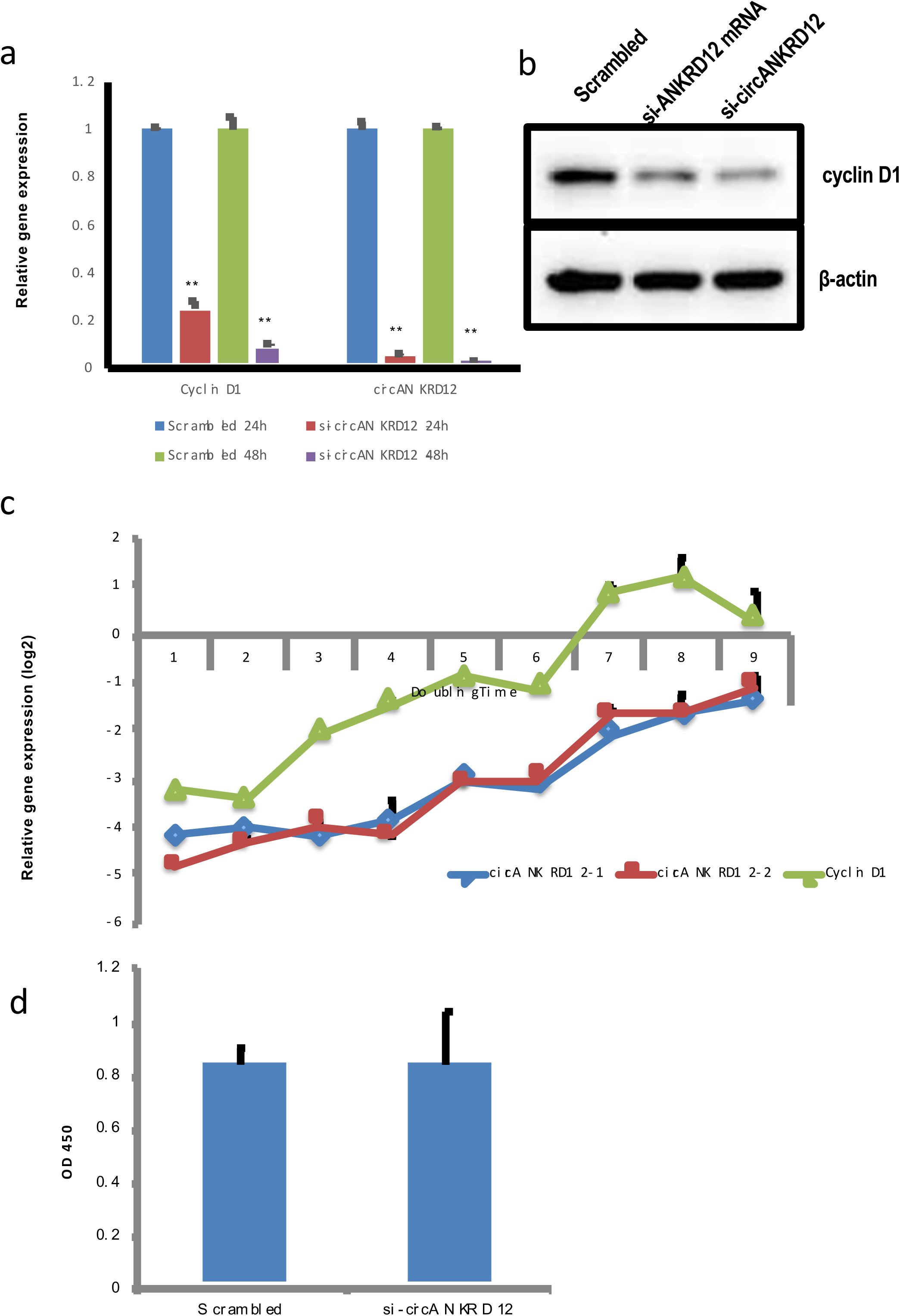
Longevity of circANKRD12 silencing in SKOV3 cells. **(a**) qRT-PCR analysis of circANKRD12 knockdown efficiency at two different time points (24h, 48h). (**b**) Western blot analysis of cyclin D1 expression in circANKRD12 and ANKRD12 silenced SKOV3 cells at 48h. (**c**) Longevity analysis of circANKRD12 silenced SKOV3 cells. qRT-PCR analysis for longevity of siRNA transfection based on knockdown efficiency of circANKRD12 in each doubling time(48h). The figure shows Knockdown efficiency of circANKRD12 on cyclin D1 expression. qRT-PCR of expression pattern of cyclin D1 from 1^st^ to 9th doubling time (d) MTS cell proliferation assay of circANKRD12 knockdown cells after 10^th^ doubling. Data in **a** is the means with error bars representing standard error of mean. \*\**P*<0.01 (Student’s *t*-test)

**Fig 8.**
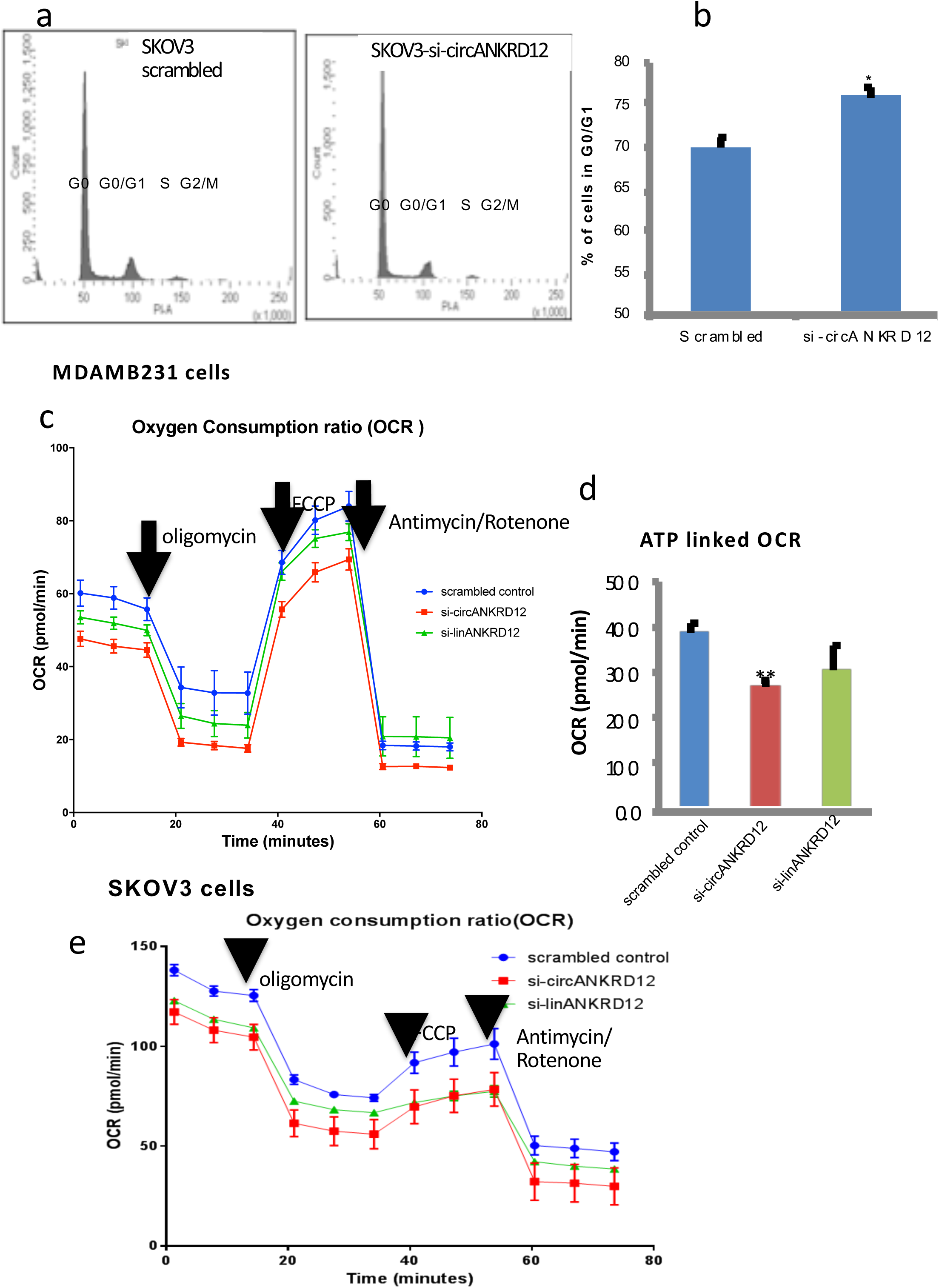
circANKRD12 silenced cells have an arrested G1stage and low OXPHOS compared to scrambled control. (**a**) Cell cycle analysis of SKOV3 cells silenced with circANKRD12 at 24h. (**b**) Bar chart representation of % of cells in G0/G1 stage. Error bars represent standard error of mean from three experimental replicates. (**c,d**) Oxygen consumption ratio and ATP production analyzed by seahorse extracellular flux analyzer in MDAMB231 cells silenced with si-circANKRD12, si-linear ANKRD12 and its control. (**e**) Oxygen consumption ratio analyzed by seahorse by extra cellular flux analysis in SKOV3 cells. Data in **c, d** and **e** are normalized with total protein(ug/ul).

### 3.8. Silencing of circANKRD12 affects Oxygen Consumption Ratio (OCR) in SKOV3 and MDAMB231 cells

AMPK signaling and other metabolic pathways are affected in circANKRD12 knockdown cancer cells (Supplementary Table 4). In order to determine whether the altered gene expression of these pathways translates to changes in basic metabolism, we analyzed metabolic phenotypes of circANKRD12 silenced MDA-MB-231 and SKOV3 cells using Seahorse extracellular flux analyzer. The oxygen consumption ratio (OCR) and ATP linked OCR analysis shows that knockdown of circANKRD12 decreases oxidative phosphorylation (OXPHOS) of MDAMB231 cells and SKOV3 cells (Fig 8 c-e). Previous reports have suggested that high invasive potential of cancer cells is negatively correlated to high energetic cancer phenotype [27,28]. These results thus indicate that circANKRD12 silencing can induce phenotypic switching between highly proliferative cells to highly invasive cells through shifting the oncobioenergetics to a low energy phenotype to facilitate invasion.

## DISCUSSION

Circular RNAs are attracting greater attention in RNA biology as there is growing evidence for their role in gene expression regulation. Even though a large repertoire of circRNAs has been identified in different organisms, tissues, diseases and developmental conditions, only a few have been evaluated for their role in cellular functions [29–32]. Functional screening using siRNA targeting is difficult for circRNAs compared to other RNA types as the choice of target region is limited to a few base pairs with specificity only at the backsplice junction. This constraint severely restricts the scale of circRNAs suitable for further functional studies using siRNA-mediated knockdown approaches.

In this study, we identified and characterized circular RNA isoforms from ANKRD12 gene (circANKRD12) that are abundantly expressed in ovarian and breast cancer cells. We showed that circANKRD12 is a stable circularized transcript, resistant to RNase R digestion and devoid of a polyA tail. In contrast to the linear mRNA form, which is predominantly nuclear, circANKRD12 is localized in the cytoplasm Differential gene expression analysis by RNA-seq of circANKRD12 silenced cells revealed that silencing may accelerate cancer cell invasion, migration and cellular movement by regulating a cascade of genes involved in these processes (supplementary Table 3, 5). Interferon signaling and cell cycle checkpoint regulation (G1/S) transition are top networks deregulated in cirANKRD12-silenced cells Cell proliferation was significantly arrested without any change in cell viability in circANKRD12 silenced SKOV3 cells. Cyclin D1, a consistently affected gene by circANKRD12 silencing, is down regulated thereby reducing cell proliferation and increasing invasion. Previous studies have reported a reduction in cyclin D1 levels and G0/G1 cell cycle stage arrest leads to an increase in migratory activities of MDA-MB-231 breast cancer cells [33]. Consistent with these findings, our results of cell cycle analysis also show a substantial G0/G1 arrest with increased migration and invasion in circANKRD12 knockdown cells. The migration and invasion assays significantly correlate with gene expression analysis and indicate an augmented cell migration and invasion rate. The 3D anchorage independent organotypic tumor models of SKOV3 show similar patterns of phenotypic characterization where invasion through collagen is increased upon silencing of circANKRD12. There is a strong phenotypic alteration after silencing circANKRD12 in the ovarian cancer cells in both 2D and 3D culture conditions suggesting that circANKRD12 is important in regulating proliferation, invasion and migration.

The Ankyrin Repeat Domain family of genes can act as putative tumor suppressors *via* p53 mediated feedback or through recruiting histone deacetylases (HDACs) to the p160 coactivator to repress transcriptional activities [17,18,34]. A clinical study of gene expression of ANKRD12 in colorectal cancer revealed that low ANKRD12 expression is correlated with overall poor survival and liver metastasis of CRC patients [19].

We observed that the silencing of ANKRD12 mRNA reduces cell proliferation, induces cell death and down regulates cyclin D1. On the contrary, silencing of circANKRD12 arrests the cell cycle progression and increases tumor invasion without significantly affecting cell viability. The knockdown of cirANKRD12 can change the oncobioenegetics as its shift from a higher OXPHOS to a lower OXPHOPS phenotype which is highly invasive. circANKRD12 may act as competing endogenous RNA (ceRNA) to regulate a circRNA-miRNA-mRNA network. Our *insilco* analysis confirms that cyclin D1 and circANKRD12 have shared binding sites for several different microRNAs [Supplementary Table 6]. Our preliminary analysis shows overexpression of hsa-miR-4768-5p reduces the level of Cyclin D1 by 20% (hsa-miR-4768-5p has common binding sites for circANKRD12 and Cyclin D1) (Data not shown). Thus, circANKRD12 could act as a microRNA sponge to regulate cyclin D1 levels.

In conclusion, our study provides the molecular, phenotypic and metabolic characterization of one of the most abundant circRNA in human ovarian and breast cancer cells. Our results suggest that circANKRD12 could be involved in a diverse set of functions ranging from cell cycle arrest, tumor invasion to immune modulation. Manipulating the levels of circANKRD12 can regulate molecular functions by altering different signaling pathways and modifies the phenotype of the cells. The distinctive change from a proliferative to a more invasive phenotype by altering cirANKRD12 levels could lead to a future circRNA based therapeutic intervention in cancer.

## ACKNOWLEDGEMENTS

The work was supported by grants from Basic Medical Research Program (BMRP) grant from Qatar Foundation to WCM-Q.

We thank Dr. Anna Halama from Dr. Karsten Suhre’s Bioinformatics Core lab at WCM-Q for providing the LL24, NCI-H226, T47D, HCC2935 cells. We also thank Ms. Aleksandra M. Liberska from Microscopy Core at WCM-Q for helping with Flow Cytometry.

